# Extracellular vesicles containing ACE2 efficiently prevent infection by SARS-CoV-2 Spike protein-containing virus

**DOI:** 10.1101/2020.07.08.193672

**Authors:** Federico Cocozza, Ester Piovesana, Nathalie Névo, Xavier Lahaye, Julian Buchrieser, Olivier Schwartz, Nicolas Manel, Mercedes Tkach, Clotilde Théry, Lorena Martin-Jaular

## Abstract

SARS-CoV-2 entry is mediated by binding of the spike protein (S) to the surface receptor ACE2 and subsequent priming by TMPRRS2 allowing membrane fusion. Here, we produced extracellular vesicles (EVs) exposing ACE2 and demonstrate that ACE2-EVs are efficient decoys for SARS-CoV-2 S protein-containing lentivirus. Reduction of infectivity positively correlates with the level of ACE2, is 500 to 1500 times more efficient than with soluble ACE2 and further enhanced by the inclusion of TMPRSS2.

## MAIN

SARS-CoV-2 is the causative agent of COVID-19 infection outbreak^1^. Viral entry into host cells is mediated by the interaction of the spike (S) protein on the surface of SARS-CoV-2 with the surface receptor angiotensin-converting enzyme 2 (ACE2)^2^. After binding to ACE2, the S protein is cleaved by TMPRSS2 and becomes fusogenic thus allowing viral entry^3^. ACE2 is expressed at the surface of pneumocytes and intestinal epithelial cells which are potential target cells for infection^4^. Soluble recombinant ACE2 neutralizes SARS-CoV-2 by binding the S protein and has proven to reduce entry of SARS-CoV-2 in Vero-E6 cells and engineered human organoids ^5^. ACE2 however is synthesized as a transmembrane protein, and we postulate that ACE2 could be present on the surface of extracellular vesicles (EVs) which could result in better efficacy as decoy to capture SARS-CoV-2.

EVs are lipid bilayer enclosed structures containing transmembrane proteins, membrane associated proteins, cytosolic proteins and nucleic acids that are released into the environment by different cell types^6^. Since EVs have the same membrane orientation as the cells, they expose at their surface the extracellular domains of transmembrane proteins that can bind to short-distant or long-distant targets. By specifically binding to different proteins and protein-containing structures, EVs can act as a decoy for virus^7^ and bacterial toxins^8^, thus having a potential role as therapeutic agents.

In order to explore the hypothesis that EVs can be used as SARS-CoV-2 decoy agents we first assessed whether ACE2 can be present in EVs from two different sources: 1) cell lines derived from tissues expressing ACE2; and 2) 293FT cells overexpressing ACE2 and TMPRSS2. As cell lines naturally expressing ACE2 we used the human lung epithelial cell line Calu3 and the epithelial colorectal cell line Caco2 which are known targets for SARS-CoV2 infection^3^. Calu3 and Caco2 were cultured in medium without FBS for 24 hours and EVs were isolated from the cell conditioned medium (CCM) by size exclusion chromatography (SEC). This technique allows the separation of EVs from soluble proteins (Figure 1A, Sup Figure 1A). We collected and analyzed EV-containing fractions, soluble protein-containing fractions and intermediate fractions containing a mixture of EVs and soluble components (Figure 1A, Sup Figure 1A). Particle quantification with nanoparticle tracking analysis (NTA) confirmed that the majority of particles released by Calu3 and Caco2 cells are isolated in EV-containing fractions (Figure 1B). Importantly, these EVs contain ACE2 protein as well as known EV markers (CD63, CD81 and ADAM10) (Figure 1C). However, high amounts of soluble ACE2 are found in the intermediate and soluble fractions obtained from CCM of these cells. In addition, despite Caco2 and Calu3 express TMPRSS2, this protease is not released in EVs or soluble fractions (Figure 1C). To obtain EVs with high amounts of ACE2 and TMPRSS2 that can be used as a decoy agent, we transduced 293FT cells with lentivirus containing ACE2 alone (293FT-ACE2) or in combination with TMPRSS2 (293FT-ACE2-TMPRSS2). 293FT cells transduced with empty plasmids were used as a control (293FT-mock). The three 293FT cell lines were cultured in FBS-containing EV-depleted medium and EVs were isolated from CCM by SEC. We observed a high particle count in EVs fractions from 293FT cells (Figure 1B), coincident with the presence of CD63, CD81, Syntenin-1 and ADAM10 EV markers (Figure 1C). ACE2 is found enriched in EVs from ACE2-transduced 293FT cells when compared to soluble fractions. Importantly, EVs from 293FT-mock and 293FT-ACE2 cells contain the cleaved form of TMPRSS2 whereas EVs from 293FT cells overexpressing TMPRSS2 also contain the full protein and its glycosylated form (Figure 1C)^9^. We detected also some particles in intermediate and soluble fractions from the three 293FT cell lines that are probably from the depleted medium and that do not contain EV markers by WB (Sup Figure 1B,1C).

**Figure 1.**
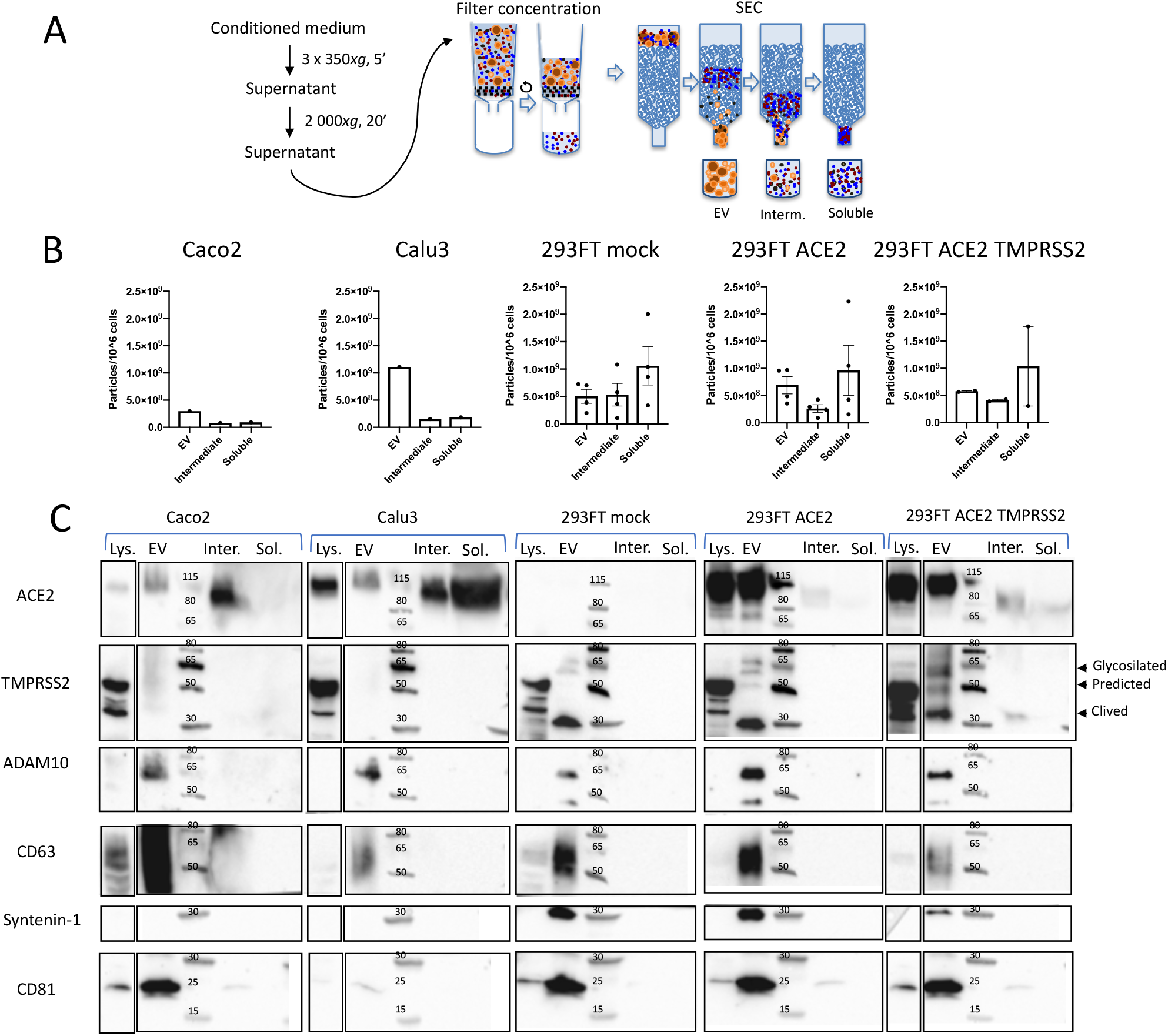
Isolation and characterization of EVs containing ACE2 and TMPRSS2. (A) Scheme of EVs isolation and separation from soluble components by SEC. (B) NTA quantification of the particles produced by 10^6^ cells contained in each fraction for different independent isolations. Error bars indicates SEM. (C) Western blot analysis of ACE2, TMPRSS2 and different EV markers in SEC fractions obtained from the five cell lines. Lysates from 4×10^5^ cells, EVs corresponding to 0,5-1×10^10^ particles and intermediate and soluble fractions from the same number of producing cells (5-34×10^6^ cells) as the EV (for Caco2 and Calu3) were loaded on the gels. Intermediate and soluble fractions from 1/10 and 1/20 producing cells (15-25×10^6^ cells), respectively, were loaded for 293FT-mock, 293FT-ACE2 and 293FT-ACE2-TMPRSS2.

We then analyzed the capacity of ACE2- and ACE2-TMPRSS2-containing EVs to reduce the infection of target cells by a lentivirus containing SARS-CoV-2-S protein. First, we determined the infectivity of the target cells Caco2, Calu3 and 293FT-ACE2 by SARS-CoV-2-S-pseudotyped lentivirus and observed that all these cell lines are infected similarly in a concentration dependent manner (Figure 2A). To assess the ability of ACE2-containing EVs to decrease virus infectivity *in vitro*, we pre-incubated SARS-CoV-2-S-pseudotyped virus with EVs isolated from 293FT-mock (MOCK-EVs) or 293FT-ACE2 (ACE2-EVs) or 293FT-ACE2-TMPRSS2 cells (ACE2-TMPRSS2-EVs) prior to the infection of target cells (Figure 2B). Infection of 293FT-ACE2 cells in the presence of ACE2-EVs and ACE2-TMPRSS2-EVs was reduced while infection remained unaffected by MOCK-EVs (Figure 2C and quantification in 2D). Importantly, this inhibition was dependent on the dose of EVs. In addition to the effect of EVs on the infection of 293FT-ACE2, Caco2 infection was also reduced in the presence of ACE2-EVs and ACE2-TMPRSS2-EVs (Figure 2E). We then quantified by ELISA the amount of ACE2 released by these cell lines. We observed that 293FT-ACE2 cells release high levels of ACE2 that is associated to EVs while 293FT-ACE2-TMPRSS2 cells release lower ACE2 levels that are equally distributed between EV and soluble fractions (Figure 2F). Strikingly, ACE2 in the soluble fractions from these latter cells was inefficient to inhibit SARS-CoV-2-S-pseudotyped virus infection as compared to the same amount of ACE2 associated to EVs (Figure 2G). Thus, considering the absolute amount of ACE2 present on EVs from these 293FT cell lines, we have observed that co-expression of the full length TMPRSS2 together with ACE2 on EVs results in a more efficient inhibition of SARS-CoV-2-S-pseudotyped viral infection (Figure 2H). Moreover, to achieve similar levels of inhibition of lentiviral infection as those observed with ACE2- or ACE2-TMPRSS2-EVs, 500 to 1500 times more of the soluble recombinant human ACE2 had to be used (Figure 2H) in accordance to previous publications^5^. Altogether, these findings highlight the increased efficiency of EVs containing full-length ACE2 to inhibit SARS-CoV-2-S-pseudotyped viral entry when compared to the soluble protein alone.

**Figure 2.**
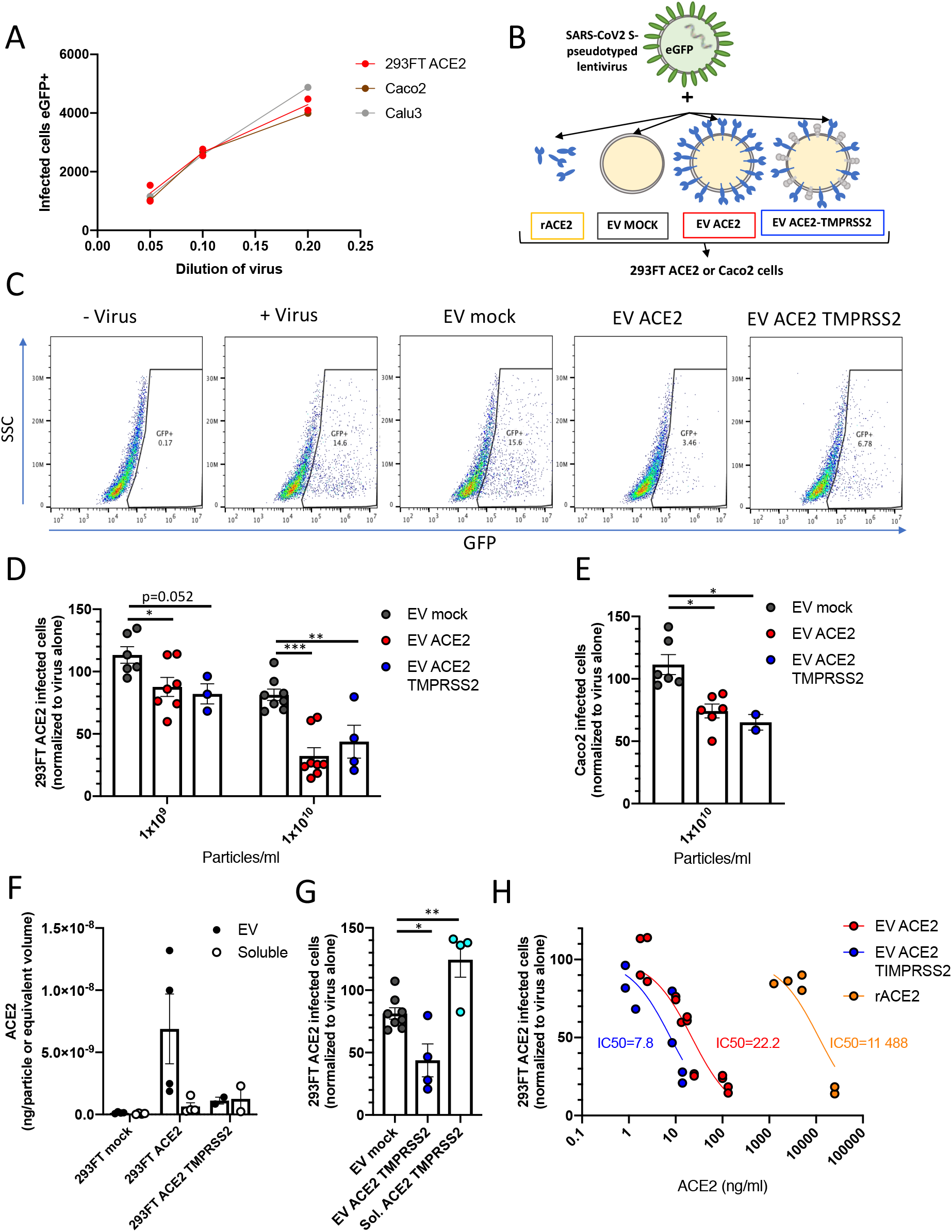
Inhibition of SARS-CoV-2-S-pseudotyped virus infection with ACE2 EVs. (A) Infection of 293FT-ACE2, Caco2 and Calu3 cells with different dilutions of a SARS-CoV-2-S-pseudotyped lentivirus encoding for eGFP. The number of infected cells was calculated by multiplying the percentage of GFP-positive cells by the initial number of cells. (B) Scheme of the infectivity assay with different pre-treatments. (C) Dot plots showing the percentage of infected 293FT-ACE2 cells obtained after incubation with viruses alone (1/10 dilution) or in combination with 1×10^10^ EV from the different 293FT cell lines. (D) Quantification of the percentage of infection of 293FT-ACE2 cells after preincubation with EVs. eGFP+ cells were measured by FACS and normalized to infection with the virus alone (100%). Results from three independent experiments are shown. All replicates from each experiment are included. *: p<0.05; **: p<0.01; (Dunnett’s test) (E) Caco2 infection in the presence of EV-ACE2 and EV-ACE2-TMPRSS2. (F) ACE2 quantification by ELISA in EV and Soluble fractions obtained from the three different 293FT cell lines. (G) Comparison of the effect on infection of EVs and soluble fractions from 293FT-ACE2-TMPRSS2. (H) Percentage of infected cells normalized to the amount of ACE2 present in EVs or as recombinant soluble form.

Our data demonstrate that EVs containing ACE2, alone or in combination with TMPRRS2, block SARS-CoV-2 Spike-dependent infection in a much more efficient manner that soluble ACE2. Thus, ACE2-EVs represent a potential versatile therapeutic tool to block not only SARS-CoV2 infection but also other coronavirus infections that use the ACE2 receptor for host cell entry, as SARS-CoV^10^and NL63^11^. The use of engineered EVs as therapeutic agents has been proposed several years ago and is currently being explored in humans^12^, proving that a well-design EV therapeutics against COVID-19 is feasible.

Note: a speculative article discussing the idea that we demonstrate experimentally here was published while we were preparing this article, thus showing concomitant emergence of similar scientific ideas^13^.

## METHODS

### Cells

Human Caco-2 (HTB-37) and Calu-3 (HTB-55) were purchased from ATCC and maintained at 37°C in a humidified atmosphere with 5% CO_2_. Caco2 and Calu3 cells were cultured in DMEM (Sigma) supplemented with 10% FBS (Gibco), 100U/ml penicillin-streptomycin (Thermo Fisher Scientific) and non-essential aminoacids (Thermo Fisher Scientific). For Calu3 cells the medium was also supplemented with 1mM sodium pyruvate (Thermo Fisher Scientific) and 10mM HEPES (Thermo Fisher Scientific). 293FT cells were cultured in DMEM medium (Sigma) supplemented with 10% FBS (Eurobio) and 100U/ml penicillin-streptomycin (Thermo Fisher Scientific). 293FT-mock, 293FT-ACE2 and 293FT-ACE2-TMPRSS2 cells were generated by stable double transduction with pTRIP-SFFV-tagBFP-2A and pTRIP-SFFV-TagRFP657-2A, pTRIP-SFFV-tagBFP-2A-hACE2 and pTRIP-SFFV-TagRFP657-2A, or pTRIP-SFFV-tagBFP-2A-hACE2 and pTRIP-SFFV-TagRFP657-2A-TMPRSS2, respectively.

### Plasmids

The plasmids psPAX2, CMV-VSVG, and pTRIP-SFFV–tagBFP-2A were previously described^14^. pTRIP-SFFV-TagRFP657-2A was generated by PCR from a synthetic gene coding for TagRFP657. pTRIP-SFFV-tagBFP-2A-hACE2 and pTRIP-SFFV-TagRFP657-2A-TMPRSS2 constructs were obtained by PCR from pLenti6-hACE2-BSD (hACE2 sequence from Addgene #1786 subcloned into pLenti6-BSD) and pCSDest-TMPRSS2 (Addgene #53887) respectively. A codon optimized version of the SARS-Cov-2 S gene (GenBank: QHD43416.1), was transferred into the phCMV backbone (GenBank: AJ318514), by replacing the VSV-G gene (phCMV-SARS-CoV-2-Spike)^15^. phCMV-SARS-CoV-2-S-H2 was obtained by PCR from phCMV-SARS-CoV-2-Spike in order to include the membrane-proximal region of the cytoplasmic domain of HIV-1 gp160 (NRVRQGYS, amino acid sequence)^16^ after residue 1246 of the S protein^17^.

### Preparation of EV-depleted Medium

EV-depleted medium was obtained by overnight ultracentrifugation of DMEM supplemented with 20% FBS at 100,000xg in a Type 45 Ti rotor (Beckman Coulter, K-factor 1042.2). After ultracentrifugation, EV-depleted supernatant was carefully pipetted from the top and leaving 7 ml in the bottom. Supernatant was filtered through a 0.22 μm bottle filter (Millipore) and additional DMEM and antibiotics were added to prepare complete medium (10% EV-depleted FBS medium).

### EV isolation by Size-Exclusion Chromatography (SEC)

239FT-mock, 293FT-ACE2 and 293FT-ACE2-TMPRSS2 cells were cultured in serum EV-depleted medium for 24h. Caco2 and Calu3 cells were cultured in FBS-free DMEM for 24h. Conditioned medium (CM) was harvested by pelleting cells at 350xg for 5 min at 4°C three times. Supernatant was centrifuged at 2,000xg for 20 min at 4°C to discard 2K pellet and concentrated on a Millipore Filter (MWCO = 10 kDa, UCF701008) to obtain concentrated conditioned medium (CCM). Medium was concentrated from 12-41 ml for Caco2 and Calu3 and from 75 ml from 293FT cells to 1 ml and overlaid on a 70nm qEV size-exclusion column (Izon, SP1). 0.5 ml fractions were collected and EVs were recovered in fractions 7 to 11 following manufacturer’s instructions. We additionally collected intermediate fractions 12 to 16 and soluble factors in fractions 17 to 21, as we previously did to analyse AChE^18^ Samples were additionally concentrated using 10kDa filter (Amicon, UCF801024) to reach a final volume of 100 μl. Samples were stored at −80°C.

### Nanoparticle Tracking Analysis (NTA)

NTA was performed to analyze EV fractions, intermediate fractions and soluble fractions using ZetaView PMX-120 (Particle Metrix) with software version 8.04.02. The instrument was set a 22°C, sensitivity 77 and shutter of 70. Measurements were done using two different dilutions, at 11 different positions (3 cycles per position) and frame rate of 30 frames per second.

### Western Blotting (WB)

Cell lysate was prepared using lysis buffer (50mM Tris, 150mM NaCl, 1% Triton, pH=8) supplemented with Phosphatase Inhibitor Cocktail (Sigma) at a concentration of 4×10^6^ cells in 100 μL of buffer. After incubation for 20 min on ice, samples were centrifuged at 18,500×g for 15 min. The pellet was discarded and the supernatant was kept for further analysis. EVs and the other SEC fractions were resuspended in 1X Laemmli Sample Buffer (Biorad) and loaded in 4-15% Mini-Protean TGX Stain-Free gels (Biorad), under non-reducing conditions. Transferred membranes (Immuno-Blot PVDF Biorad) were developed using Clarity Western ECL substrate (Biorad) and the ChemiDoc Touch imager (Biorad). Antibodies for WB were anti-human: ACE2 (clone EPR4435, Abcam 108252), TMPRSS2 (clone EPR3681, Abacam 92323), ADAM10 (clone 163003, R&D Systems MAB1427), CD63 (clone H5C6, BD Bioscience 557305), Syntenin-1 (clone C2C3, Genetex GTX10847) and CD81 (clone 5A6, Santa Cruz sc-23692). Secondary antibodies included HRP-conjugated goat anti-rabbit IgG (H+L) (Jakson 111-035-144), HRP-conjugated goat anti-muse IgG (H+L) (Jakson 111-035-146).

### Viral Production

SARS-CoV-2-S-pseudotyped lentiviruses were produced by transient transfection of 293FT cells in 150 cm^2^ flasks with 5 μg phCMV-SARS-Cov-2-S-H2, 13 μg psPAX2 and 20 μg pTRIP-SFFV-eGFP-NLS and 114 ul of TransIT-293 (Mirus Bio). One day after transfection, media was removed and fresh media was added. SARS-CoV-2-S-pseudotyped viruses supernatant was centrifuged at 300xg for 10 min to remove dead cells, filtered with a 0.45 μm filter (Millipore) and loaded on top of a 20% sucrose gradient for concentration. Viral concentration was achieved by ultracentrifugation at 120,000xg for 1h 30 min in a SW32i rotor. The pellet containing concentrated SARS-CoV-2 S-pseudotyped virus was resuspended in 1 ml depleted DMEM and 100 μl aliquots were stored at −80°C.

### Infectivity Assay

10,000-20,000 293FT-ACE2, Caco2 and Calu3 cells were seeded in a 96 well plate and after 6 h infected with SARS-CoV-2 S-pseudotyped virus in EV-depleted medium. Infection was performed in the absence or in the presence of different amount of EVs or human recombinant ACE2 (Abcam, 151852). Cells were then spinoculated at 1,200xg for 1h 30 min at 25°C. 48h after infection, cells were trypsinized, fixed and infection was measured by analyzing eGFP expression using a CytoflexLX cytometer. Data was analyzed using FlowJo software.

### ACE2 Enzyme-Linked Immunosorbent Assay (ELISA)

Quantification of the amount of human ACE2 in the different EV and fractions was done using the human ACE2 ELISA kit (Abcam, ab235649) following manufacturer’s instructions.

## ACKNOWLEDGEMENTS

This work was supported by Institut Curie, INSERM, CNRS, grants H2020-MSCA-ITN (722148, TRAIN-EV), INCa (11548) and Fondation ARC (PGA1 RF20180206962) to C Théry, LABEX DCBIOL (ANR-10-IDEX-0001-02 PSL* and ANR-11-LABX-0043) to C. Théry and N. Manel, LABEX VRI (ANR-10-LABX-77), ANRS (France Re-cherche Nord & Sud Sida-hiv Hépatites; ECTZ36691, ECTZ71745), Sidaction (17-1-AAE-11097-2), ANR (ANR-19-CE15-0018-01, ANR-18-CE92-0022-01), DIM1HEALTH to N Manel.

## AUTHOR CONTRIBUTIONS

FC, EP, NN, XL performed the experiments. FC, EP, MT, LMJ, CT analyzed the data. LMJ, CT, MT designed the experiments. EP, FC, MT, CT, LMJ wrote the paper. XL, NM and JB, OS designed plasmids, XL, NM generated cells overexpressing ACE2 and TMPRSS2 and developed the infection assay.

## Supplementary

**Supplementary Figure.**
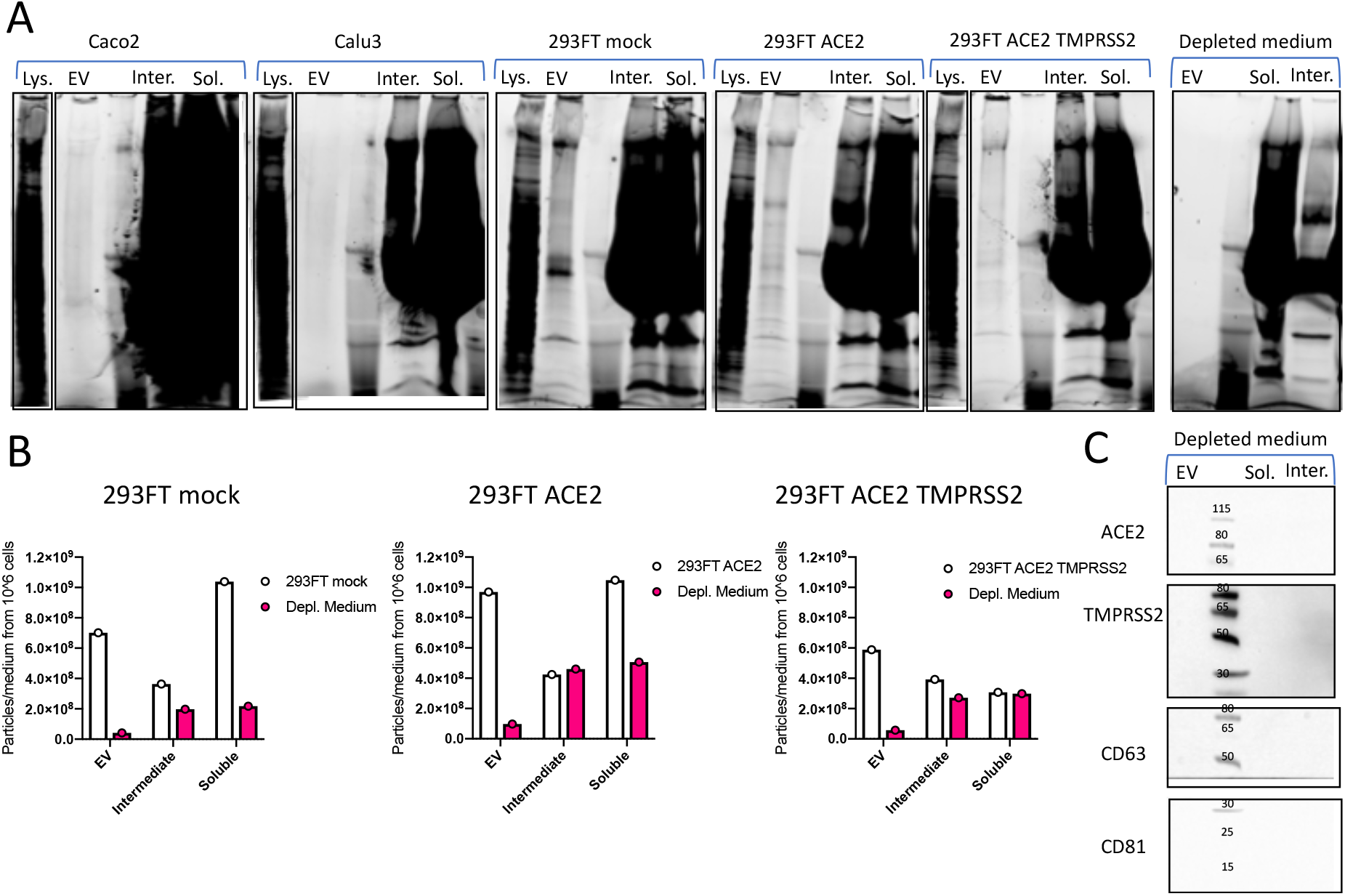
Depleted medium particle contribution to the different fractions isolated by SEC. (A) Protein stain-free images of gels used for WB in Figure 1C. (B) Number of particles counted in each fraction of 293FT-mock, 293FT-ACE2 and 293FT-ACE2-TMPRRS2 cell CCM, compared with non-conditioned depleted medium isolation performed in parallel. Numbers of particles are normalized to the volume of medium used for each purification. (C) Western blot analysis of nonconditioned depleted medium fractions isolated by SEC (material loaded on the WB was obtained from 15 ml initial volume of depleted medium for EV, 1,5 ml for intermediate and 0,75 ml for soluble).

## REFERENCES

1. Zhou, P. et al. A pneumonia outbreak associated with a new coronavirus of probable bat origin. Nature 579, 270–273 (2020).

2. Walls, A. C. et al. Structure, Function, and Antigenicity of the SARS-CoV-2 Spike Glycoprotein. Cell 181, 281–292.e6 (2020).

3. Hoffmann, M. et al. SARS-CoV-2 Cell Entry Depends on ACE2 and TMPRSS2 and Is Blocked by a Clinically Proven Protease Inhibitor. Cell 181, 271–280.e8 (2020).

4. Ziegler, C. G. K. et al. SARS-CoV-2 Receptor ACE2 Is an Interferon-Stimulated Gene in Human Airway Epithelial Cells and Is Detected in Specific Cell Subsets across Tissues. Cell 181, 1016–1035.e19 (2020).

5. Monteil, V. et al. Inhibition of SARS-CoV-2 Infections in Engineered Human Tissues Using Clinical-Grade Soluble Human ACE2. Cell 181, 905–913.e7 (2020).

6. Mathieu, M., Martin-Jaular, L., Lavieu, G. & Théry, C. Specificities of secretion and uptake of exosomes and other extracellular vesicles for cell-to-cell communication. Nat. Cell Biol. 21, 9–17 (2019).

7. de Carvalho, J. V et al. Nef Neutralizes the Ability of Exosomes from CD4+ T Cells to Act as Decoys during HIV-1 Infection. PLoS One 9, e113691 (2014).

8. Keller, M. D. et al. Decoy exosomes provide protection against bacterial toxins. Nature 579, 260–264 (2020).

9. Afar, D. E. H. et al. Cancer Research. Cancer Res. 59, 6015–6022 (2001).

10. Li, W. et al. Angiotensin-converting enzyme 2 is a functional receptor for the SARS coronavirus. Nature 426, 450–454 (2003).

11. H, H. et al. Human Coronavirus NL63 Employs the Severe Acute Respiratory Syndrome Coronavirus Receptor for Cellular Entry. Proc. Natl. Acad. Sci. U. S. A. 102, (2005).

12. Wiklander, O. P. B., Brennan, M. Á., Lötvall, J., Breakefield, X. O. & Andaloussi, S. EL. Advances in therapeutic applications of extracellular vesicles. Sci. Transl. Med. 11, (2019).

13. Inal, J. M. Decoy ACE2-expressing extracellular vesicles that competitively bind SARS-CoV-2 as a possible COVID-19 therapy. Clin. Sci. 134, 1301–1304 (2020).

14. Cerboni, S. et al. Intrinsic antiproliferative activity of the innate sensor STING in T lymphocytes. J. Exp. Med. 214, 1769–1785 (2017).

15. Grzelak, L. et al. SARS-CoV-2 serological analysis of COVID-19 hospitalized patients, pauci-symptomatic individuals and blood donors. medRxiv 2020.04.21.20068858 (2020). doi: 10.1101/2020.04.21.20068858

16. F, M., E, K., J, S., A, B. & HG, G. Rescue of human immunodeficiency virus type 1 matrix protein mutants by envelope glycoproteins with short cytoplasmic domains. J. Virol. 69, 3824–3830 (1995).

17. Moore, M. J. et al. Retroviruses Pseudotyped with the Severe Acute Respiratory Syndrome Coronavirus Spike Protein Efficiently Infect Cells Expressing Angiotensin-Converting Enzyme 2. J. Virol. 78, 10628–10635 (2004).

18. Liao, Z. et al. Acetylcholinesterase is not a generic marker of extracellular vesicles. J. Extracell. Vesicles 8, (2019).

